# High precision animal stereotaxis with 3D polychromic photogrammetry

**DOI:** 10.64898/2026.07.27.741054

**Authors:** Achim Klug, Matthew Ridenour, Benzheng Li, Jason Jacoby, Adam Dau, Tim C. Lei

## Abstract

Stereotaxic brain surgery is a foundational neurosurgical technique used to deliver chemical, pharmacological, or genetic material to specific brain regions, or to precisely implant electrodes and stimulators. Accurate targeting depends on reliable identification of skull landmarks, particularly bregma and lambda. Here we present a photogrammetry-based automated small animal stereotaxic platform that uses a single, freely handheld camera, such as a standard smartphone, to generate high-resolution, polychromatic 3D skull reconstructions. Because photogrammetry requires no fixed overhead hardware, the surgical field remains fully accessible for instruments, microscopes, and other equipment. The resulting color reconstructions substantially improve identification of bregma and lambda compared to monochromatic approaches, and the higher spatial resolution translates directly into improved targeting accuracy and surgical speed. The system operates by a user moving a handheld camera around the exposed skull. The platform automatically produces a detailed 3D mesh and computes stereotaxic coordinates without manual measurement. Together, these properties yield a practical, accessible yet accurate platform for automated small animal neurosurgery. We previously described a structured illumination approach in combination with a Steward platform; however, that system required a fixed projector and camera array that occupied critical surgical workspace and produced monochromatic meshes that complicated landmark identification. The photogrammetry-based method described here overcomes both limitations while retaining full compatibility with the Steward platform as a stereotaxic base.

## 1. Introduction

Stereotaxic brain surgery is a foundational neurosurgical technique that enables precise, targeted delivery of chemical, pharmacological, or genetic material to discrete brain regions, as well as accurate implantation of electrodes and other devices (Horsely and Clark, 1908, Cetin et al. 2006). The procedure typically requires the surgeon to first align the subject’s skull into the so-called “skull-flat” position, defined such that the line connecting the bregma and lambda landmarks plus the line connecting the two sides of the skull (red lines in fig. 1) are both parallel to the surgical plane (Paxinos and Watson, 1982). Once the skull has been leveled, the surgeon performs a craniotomy, through which a pipette, electrode, or other tool is inserted into the brain at a prescribed angle and depth to reach the intended target. Despite its central importance to neuroscience research, this fundamental technique has remained largely unchanged for decades in small animal subjects, relying on imprecise mechanical measurement or purely visual alignment (“eyeballing”) to establish the skull-flat position. Such imprecision is particularly consequential when targeting deep brain nuclei, where even small positioning errors translate into longer surgeries, degraded targeting accuracy, and, in many cases, outright procedural failure (Rangarajan et al., 2016).

**Figure 1.**
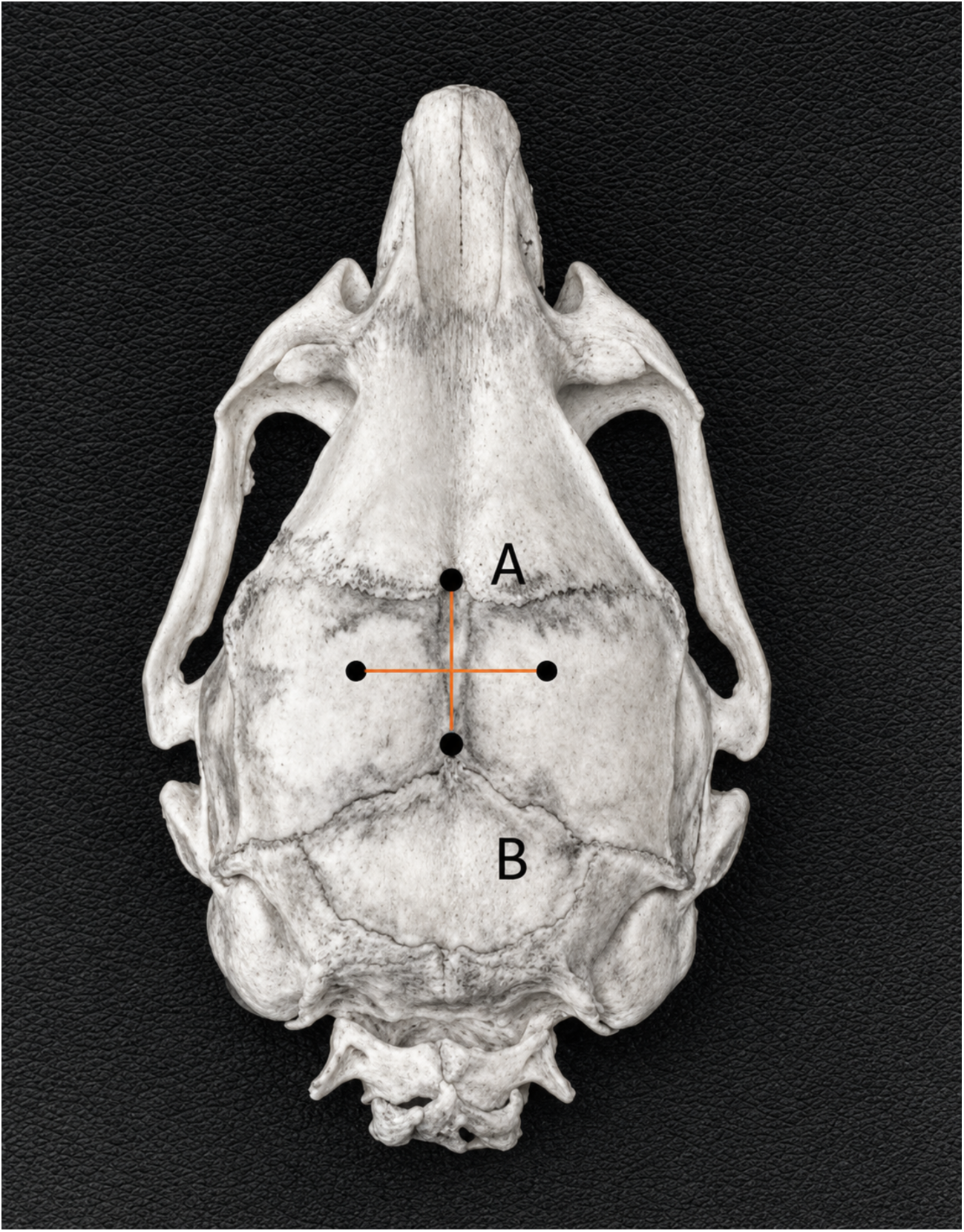
Sketch of a generic rodent skull, showing bone sutures and landmarks. The bregma (A) landmark, defined by the intersection of the sagittal and coronal sutures, and lambda (B) defined by the intersection of the sagittal and lambdoid sutures.

Over the past several years, small animal stereotaxic neurosurgery has begun to move away from manually operated, vernier-read frames toward electronically guided and robotic platforms designed to reduce operator-dependent variability. The Angle Two and Angle Three stereotaxic instruments (Leica Biosystems, Nussloch, Germany) introduced encoder-equipped manipulators linked to atlas-referenced software that perform bregma-lambda scaling and skull-tilt correction (Leica Biosytems, n.d.). Neurostar (Tübingen, Germany) developed a Drill and Injection Robot that performs craniotomies (Neurostar, n.d.), and the Model 71000 stereotaxic apparatus (RWD Life Science, Schenzhen, China) couples a built-in rodent brain atlas with automated craniotomy and injection steps that are conventionally performed by hand (RWD Life Science Co., Ltd., n.d.). Yet even these automated platforms continue to rely on point-based landmark measurement or atlas-only registration, leaving accurate, rapid targeting of small, deep, and anatomically variable structures, such as brainstem nuclei, an unresolved challenge.

Here, we address this gap by combining three-dimensional computer vision, specifically color-resolved (polychromatic) photogrammetry (Linder, 2009, Förstner and Wrobel, 2016, Szeliski, 2022,), with a six-degree-of-freedom (6DOF) robotic positioning platform to create a fully automated small animal stereotaxic system capable of accurately and rapidly targeting brain nuclei, including previously hard-to-reach targets deep within the brainstem. Photogrammetry is a computational computer vision technique in which ordinary two-dimensional photographs, captured by a freely positioned camera from multiple angles around an object (here, the exposed dorsal surface of the rodent skull), are used to compute the physical three-dimensional location (x, y, and z) of each point on the object’s surface (Hartley and Zisserman, 2004, Özyeşil et al. 2017). These computed coordinates yield a highly detailed, micrometer-resolution 3D reconstruction of the rodent skull. The native surface color at each point can also be extracted from the source images and superimposed onto the reconstructed mesh, substantially aiding identification of cranial suture landmarks such as bregma and lambda. This approach is both faster and more accurate than conventional manual methods for locating these landmarks: physical probing of the skull is replaced by a brief series of photographs, and the resulting 3D skull profile is used to compute angular deviation from the skull-flat position directly, enabling precise positional correction.

To achieve maximal targeting accuracy, the mechanical design of this stereotaxic system departs substantially from conventional practice. In traditional stereotaxic systems, the rodent is secured to an immobile platform using ear bars, bite bars, or head mounts, while a mechanical stereotaxic arm, equipped with multiple manipulators that are either manually or electronically controlled, is positioned at the desired coordinates. A stereotaxic tool, such as an injection needle or neural recording probe, is then advanced from above the rodent’s skull to the prescribed insertion point and angle (Horsely and Clark, 1908, Cetin et al. 2006). This configuration is prone to positional error: because the mechanical arm sits well above the skull, its extended, multi-segment length is susceptible to mechanical instability and vibration (figure 2A). In this work, we instead use a 6DOF platform, three translational and three rotational axes, to align the rodent’s skull to a stationary, vertically oriented insertion arm, minimizing mechanical instability and insertion error (figure 2B). This platform is adapted from the classic Stewart-platform design (Stewart, 1965) but replaces its conventional linear actuators with rotary servo motors, reducing platform height and thereby increasing mechanical stability. With this approach, the platform automatically positions the animal’s skull into the skull-flat orientation and subsequently moves it into the correct surgical position under precise electronic motor control, achieving a level of surgical speed and targeting accuracy that manual methods cannot match. Existing techniques typically require the user to measure skull landmarks manually, advancing a tool along the z-axis to record the z-coordinate of each landmark (figure 3) and adjusting skull position by hand until skull-flat is reached (Cetin et al, 2006, Kawa et al. 2024). This process is slow and imprecise, since it reduces a complex three-dimensional skull profile to only a handful of points; sampling additional points would improve accuracy, but manual measurement makes this impractical within a reasonable surgical timeframe. We therefore hypothesized that automated computer vision could sample many more points within the same timeframe, simultaneously improving both the accuracy and the speed of the procedure.

**Figure 2.**
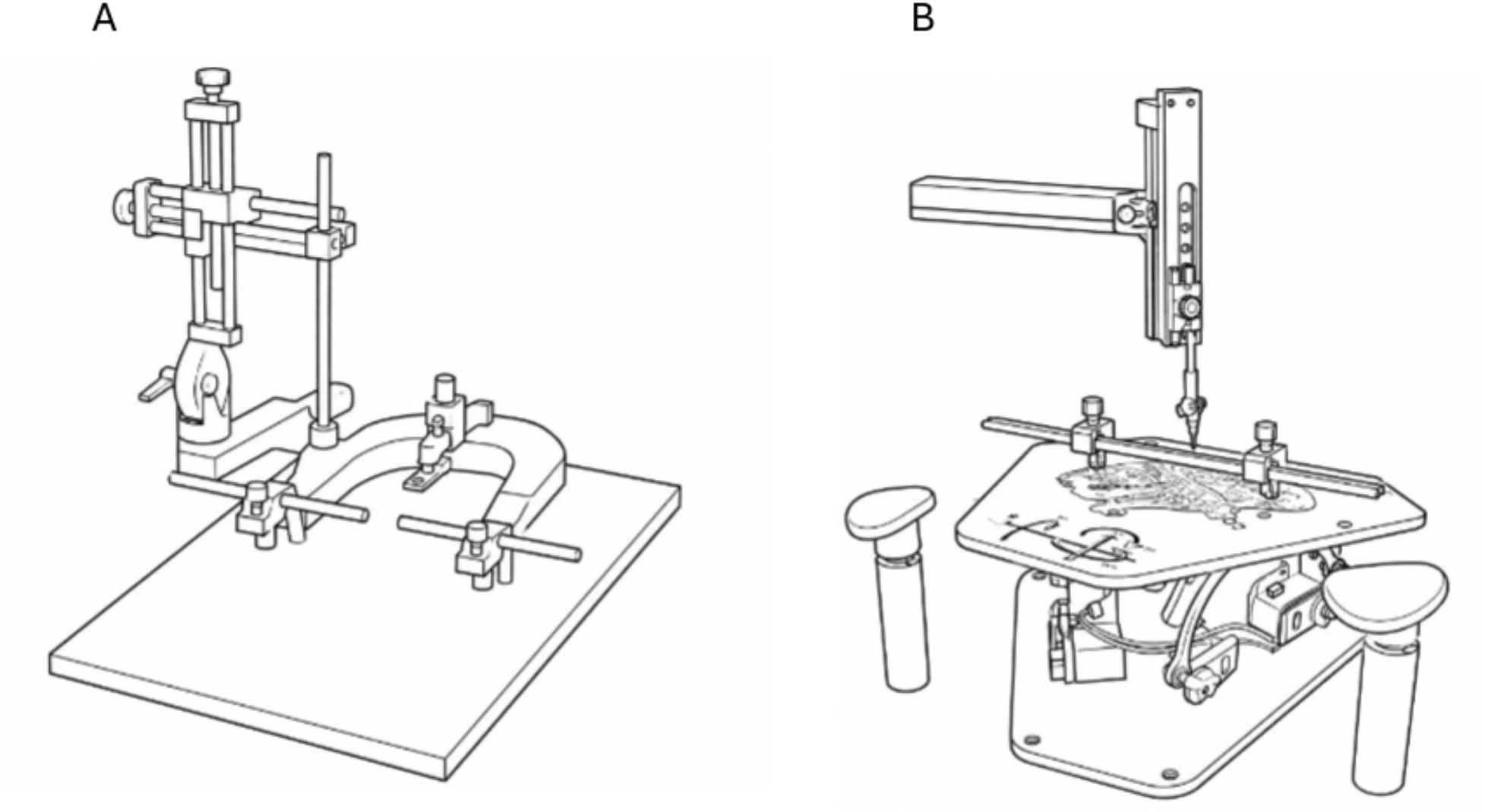
Comparison of traditional stereotaxic devices with hexapod based designs. A: In traditional stereotaxic systems, the rodent is secured to an immobile platform using ear bars, bite bars, or head mounts, while a mechanical stereotaxic arm, equipped with multiple manipulators that are either manually or electronically controlled, is positioned at the desired coordinates. B: In hexapod designs, three translational and three rotational axes align the rodent’s skull to a stationary, vertically oriented insertion arm, minimizing mechanical instability and insertion error. This platform is adapted from the classic Stewart-platform design but replaces its conventional linear actuators with rotary servo motors, reducing platform height and thereby increasing mechanical stability.

**Figure 3.**
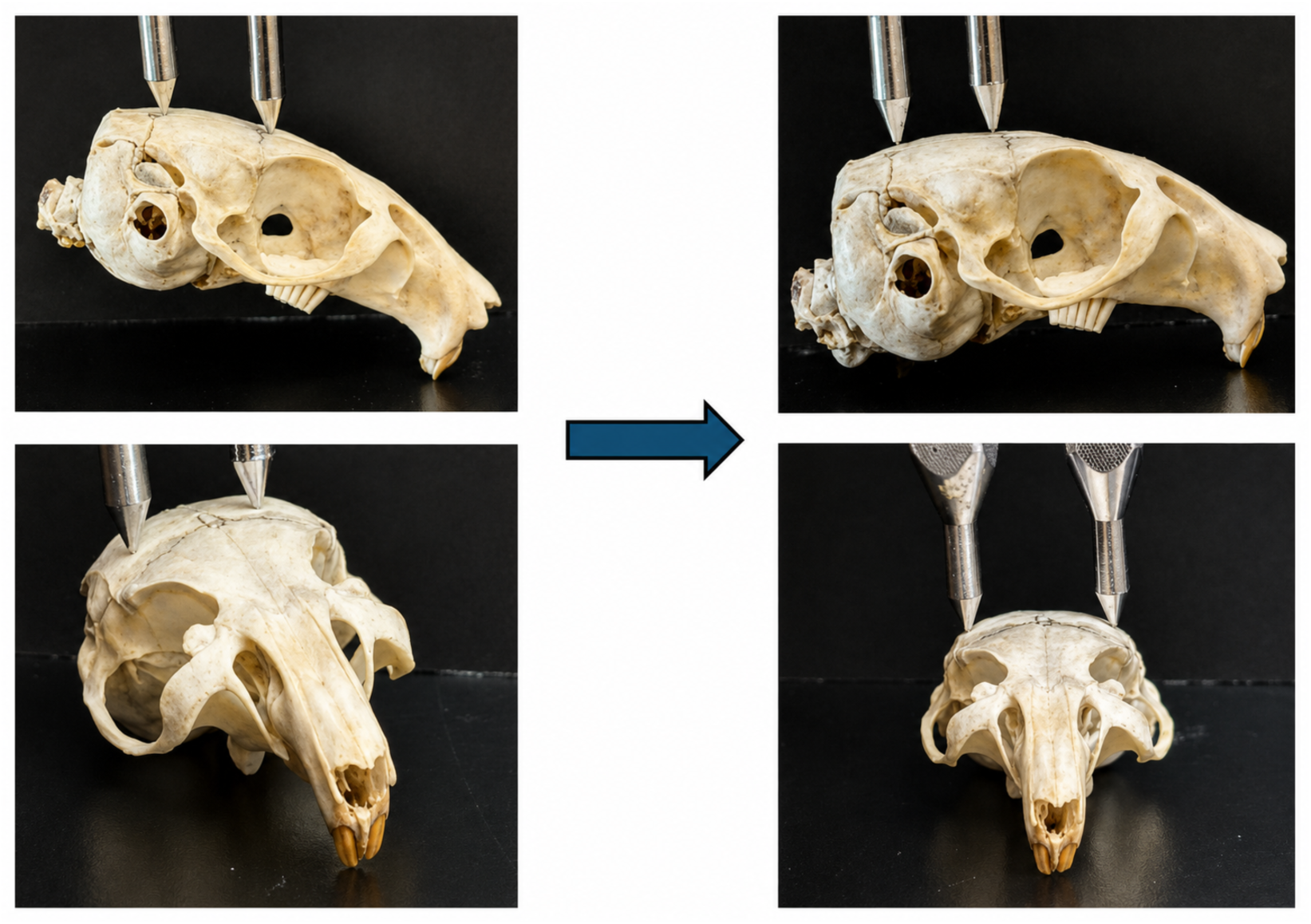
An illustration of current methods used to flatten animal skulls for stereotaxic surgery. The lambda and bregma landmarks are probed to obtain reading of their relative positions in the z-axis. The skull is then adjusted until these two points are at equal height, leveling the pitch axis. This process is then repeated with two lateral points defining a line perpendicular to the line defined by the lambda and bregma points, which levels the roll axis.

In a previous report, we described a structured-illumination subsystem (Ly et al. 2022, DePiero et al., 1996) (figure 4) and its use for 3d skull reconstruction. Structure illumination is now replaced with photogrammetry. Unlike structured illumination, photogrammetry requires neither projected optical contrast patterns nor a fixed camera array; a handheld, high-resolution mobile phone camera is sufficient to capture the images needed for reconstruction. This eliminates the need for an overhead optical projector or fixed cameras, freeing space above the animal for surgical equipment and substantially simplifying use of the technique in the laboratory setting. As described above, the resulting 3D mesh retains the native surface coloration of the skull, further simplifying identification of the bregma and lambda landmarks. The original Stewart-platform design was likewise modified to integrate fully with this photogrammetry-based workflow.

**Figure 4.**
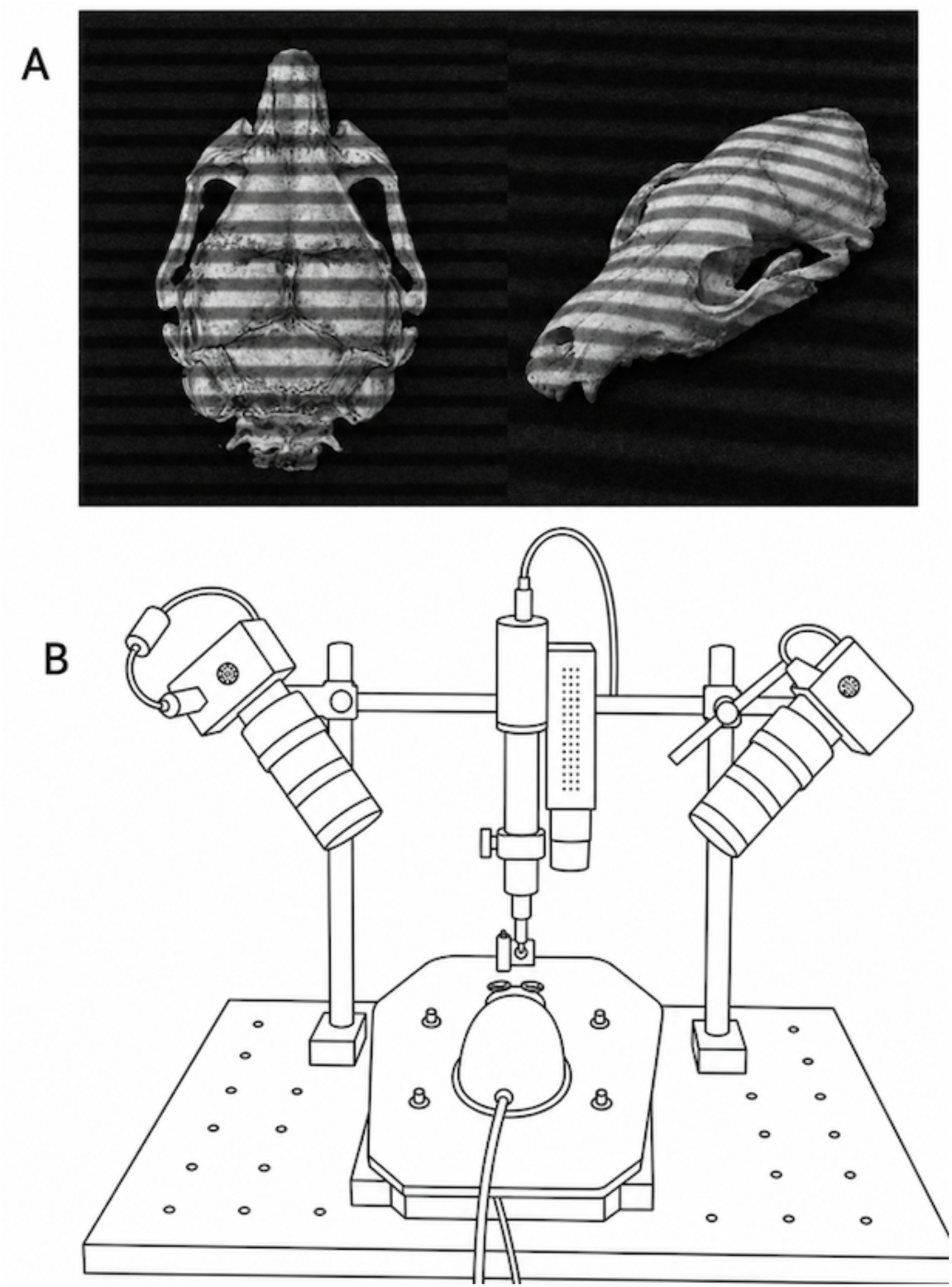
An example of the structured illumination used to approximate a 3D mesh reconstruction of skull shape in previous iterations of this system. (A) Skulls illuminated with a structured illumination pattern. (B) An schematic of a previous iteration of this system, showing a projector and cameras mounted on a gantry above the platform

## 2. Methods

In this report, photogrammetry replaced structured illumination as the method for mapping the surface of the animal skull into virtual space, generating a high-resolution, polychromatic 3D mesh. Unlike structured illumination, which requires the fixed and precise placement of an optical projector and two cameras above the skull, photogrammetry uses a single, freely positioned digital camera to capture a series of images around the skull. The primary requirements are that the images provide adequate coverage of the skull, are captured from multiple angles, and are taken at a consistent magnification (zoom setting). Approximately 20 to 30 images are sufficient to generate a 3D skull mesh with the accuracy required for stereotaxic surgeries targeting small, deep brain nuclei. Below, we briefly describe the two key algorithms underlying this process, Scale-Invariant Feature Transform (SIFT) and Multi-View Stereo (MVS), which together enable photogrammetry to generate polychromatic, highly detailed 3D meshes of the skull surface. We also describe the modifications made to the computer-controlled Stewart platform required for compatibility with photogrammetry.

### 2.1 Scale-Invariant Feature Transform (SIFT) for keypoint identification and image stitching

In photogrammetry (figure 5), images acquired from multiple angles are first stitched together by aligning keypoints, image locations with distinctive local features. Rather than relying on manual keypoint selection, the Scale-Invariant Feature Transform (SIFT) algorithm (Lowe, 1999, 2004) identifies these keypoints automatically. The algorithm then generates feature descriptors that remain invariant to translation, scaling, and rotation, and are resistant to illumination variation. These descriptors are used to match keypoints across the acquired images for stitching. Once the images are stitched, the camera positions from which they were taken can be estimated using geometric triangulation. Based on these estimated camera positions, the positions of the keypoints can also be estimated, yielding a coarse point cloud of the animal skull.

**Figure 5.**
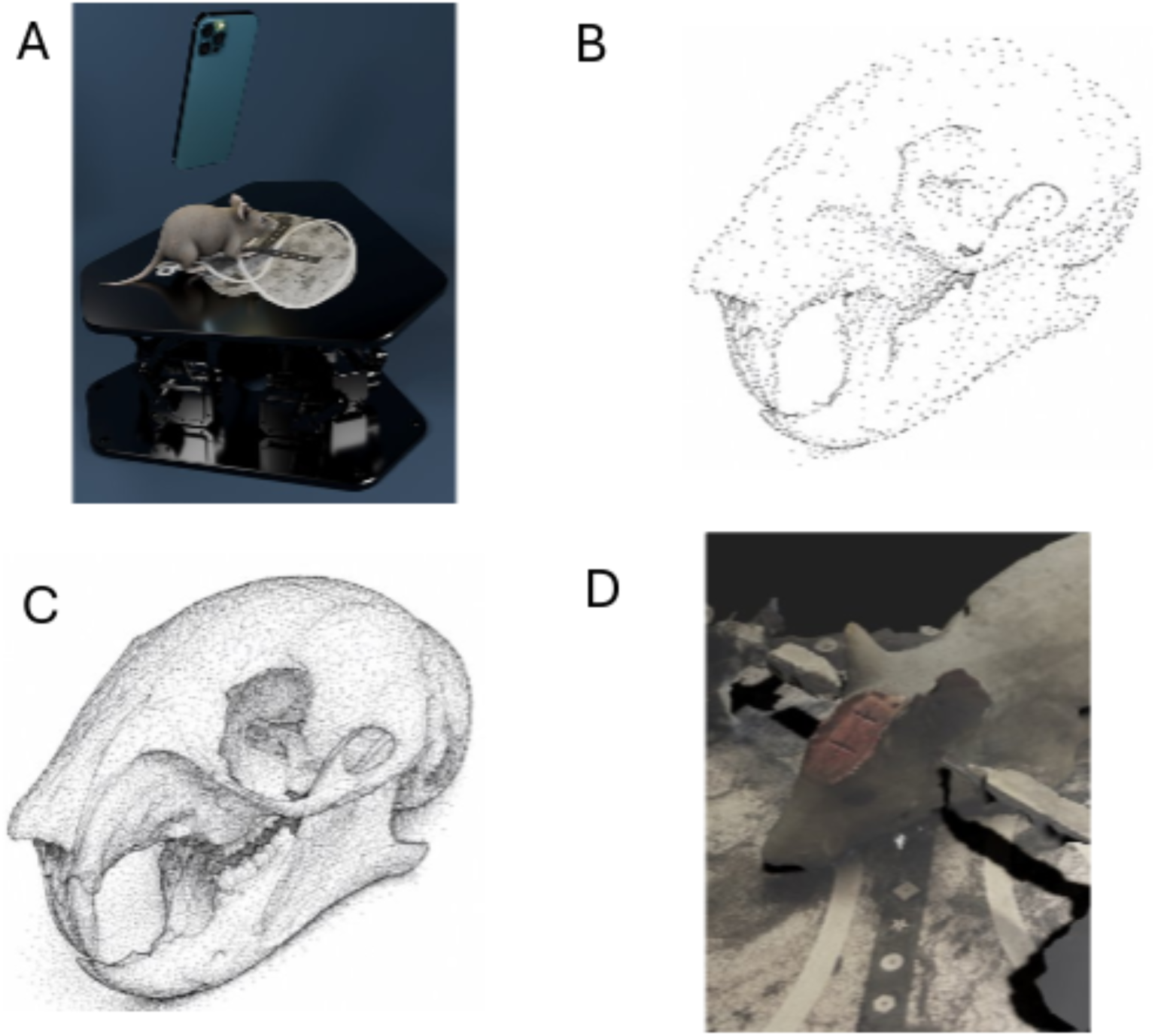
Workflow of the photogrammetry process, beginning with the acquisition of images with a cell phone camera and ending with a detailed 3D mesh reconstruction. (A) A cell phone camera is used to capture images from various angles around the subject. (B) Scale -Invariant Feature Transform (SIFT) algorithm identifies key points, stitches images, and estimates camera distance via triangulation. A rough point cloud is created. (C) Multi-view stereo (MVS) algorithm estimates more vertices and their 3D positions. (D) A detailed mesh is created and overlaid with color via a texturing step.

The SIFT algorithm begins by blurring the acquired images through convolution with a Gaussian spatial distribution, using a series of progressively increasing standard deviations (σ). Subtracting each blurred image from the original generates a corresponding difference image. Local maxima within these difference images are identified as keypoints, since they represent locations of strong local contrast relative to neighboring pixels, a property that makes them well suited as keypoints. Because the algorithm relies on intensity gradients rather than absolute intensity values, it is largely insensitive to illumination fluctuations. Rotational dependence is removed by computing the local gradient directions around each keypoint, identifying the dominant orientation, and using it to rotate the keypoint’s local image patch into a canonical orientation. Finally, scale dependence is eliminated by determining the characteristic scale at each keypoint and rescaling its local image information accordingly, rendering the keypoint description scale invariant.

Once keypoints are identified, a descriptor is calculated for each. These descriptors are two-dimensional feature vectors encoding local image gradient information, magnitude and direction, computed by comparing each keypoint to its neighboring pixels. These feature vectors uniquely characterize the local image content around each keypoint, enabling efficient matching of corresponding points across images for stitching. Standard feature-matching techniques, such as nearest-neighbor search, are then used to pair keypoints based on the similarity of their descriptor vectors.

Based on the matched keypoints between images, the camera positions and orientations from which the images were taken are estimated. This estimation relies on the fact that matched keypoints must satisfy the epipolar constraint, which specifies that for any point observed in one image, the corresponding point in a second image must lie along the epipolar line determined by the geometric relationship between the two camera positions from which the images were captured. To achieve high accuracy, camera positions are refined iteratively through global optimization until a global error minimum is reached. Initially, only the two images with the greatest number of matched keypoints are used to estimate the positions and orientations from which those images were captured. Additional images are then incorporated one at a time, with camera positions and orientations re-estimated at each step until all images have been incorporated into the reconstruction. Once camera positions and orientations are established, the positions of the keypoints in virtual 3D space can be estimated through geometric triangulation, similar to the approach used in structured illumination. However, because only a limited number of keypoints are identified, the resulting point cloud of the animal skull is sparse. Additional image-processing steps are therefore required to add further vertices to the point cloud, ensuring sufficient spatial detail for accurate stereotaxic alignment.

### 2.2 Detailed skull mesh generation with the Multi-View Stereo algorithm

Once the 3D positions of the keypoints have been determined, the Multi-View Stereo (MVS) algorithm (figure 6, Seitz & Dyer 1999, Seitz et al. 2006, Furukawa & Ponce, 2010)) is used to estimate additional vertices, including for pixels that were not identified as keypoints. The algorithm first selects two neighboring images with significant spatial overlap, then attempts to match each non-keypoint pixel in the first image to a corresponding pixel in the neighboring image, even in regions that lack strong image contrast. This matching is possible because corresponding pixels must lie along the epipolar lines defined by the geometric relationship between the two camera positions from which the images were captured. Based on this principle, the algorithm searches along the corresponding epipolar line in the neighboring image to identify the best-matching pixel. This process is repeated to estimate the 3D positions of all non-keypoint pixels in the first image, and then repeated in turn for each remaining image. Because of the many possible image combinations, this iterative process can estimate the same physical point on the skull multiple times; a merging step therefore fuses these duplicate estimates into a single, coherent, dense 3D point cloud. Filtering is also performed to eliminate outlier points, ensuring the smoothness and consistency of the surface.

**Figure 6.**
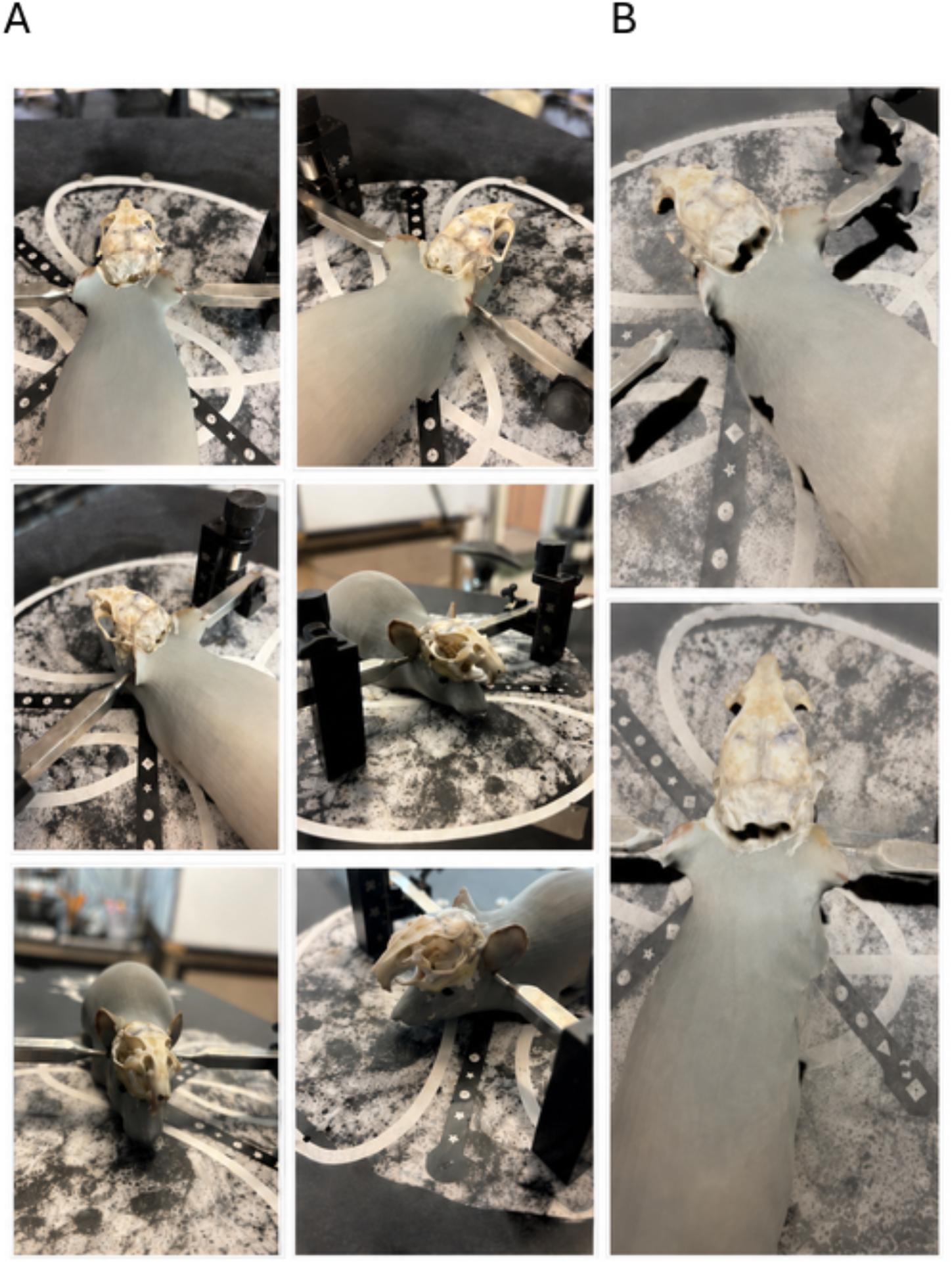
A set of single 2D images is used to compute a 3D reconstruction. (A) A subset of the images captured on a mobile phone camera used to create a 3D reconstruction, using a model mouse with a gerbil skull affixed to the head. (B) Two screenshots of the 3D reconstruction created from the full image set via photogrammetry

The MVS algorithm thus yields a dense point cloud containing the estimated 3D coordinates of the skull surface. For use in stereotaxic positioning, however, this point cloud must first be converted into a 3D mesh, which involves connecting its vertices with edges using algorithms such as Delaunay tetrahedralization. Once the mesh is generated, it is overlaid with color information through a texturing step, producing a polychromatic rendering in virtual space. This color information is extracted directly from the source 2D images, since each mesh vertex can be readily mapped back to its corresponding pixel. This color information is essential for reliably identifying skull suture landmarks, such as bregma and lambda, required for accurate stereotaxic targeting.

### 2.3 Improvements to the 6DOF Stewart platform

The original 6DOF, servo-driven Stewart platform provides a mechanically stable means of rapidly positioning the animal’s skull for stereotaxic surgery under computer control. However, several modifications were required to adapt the platform for compatibility with photogrammetry-based imaging.

Unlike structured illumination, in which the fixed positions of the projector and two cameras provide an absolute spatial reference for 3D reconstruction (Ly et al., 2021), photogrammetry allows the camera to be freely positioned and its magnification adjusted. Consequently, the translational dimensions (x, y, and z axes) of the 3D mesh generated by photogrammetry are not absolute but relative: for example, a small object photographed at high zoom can be reconstructed to appear the same size as a larger object photographed at lower zoom. It is therefore essential to modify the Stewart platform to enable accurate scaling of the translational dimensions of the reconstructed mesh.

Both issues, scaling and surface reconstruction, were addressed by engraving a high-contrast pattern, interwoven with sets of fiducial marks at predetermined intervals, onto the platform’s top surface (figure 7). Within each set, three fiducial marks define two perpendicular translational axes (x and y) on the platform surface. Multiple sets were engraved across the platform to ensure that at least one set remains unobstructed by the animal, regardless of body size, for use in scaling adjustments. As described above, the SIFT algorithm requires optical contrast to identify keypoints; because the platform surface, unlike the animal skull, lacks the natural contrast provided by sutures and other structural features, the engraved pattern supplies the artificial contrast needed to generate sufficient keypoints for a dense, high-fidelity photogrammetric reconstruction of the surface.

**Figure 7.**
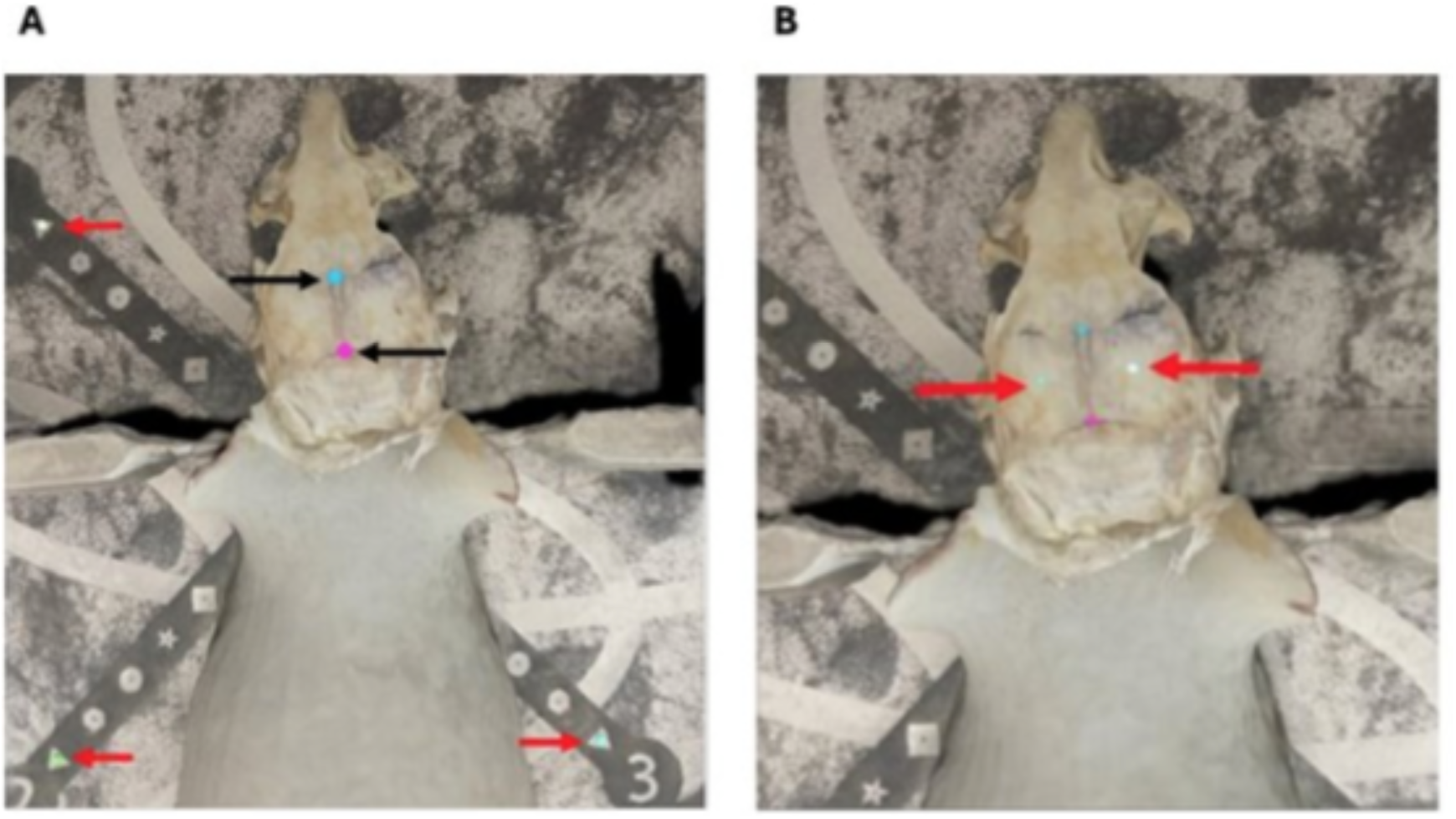
The process of 3D reconstruction. (A) A 3D reconstruction of a model mouse with a gerbil skull affixed to the top of the head. Red arrows denote fiducial markers on the base plate surface. Black arrows indicate lambda and Bregma. (b) The same model with lateral points forming a line perpendicular to the line between Lambda and Bregma calculated. These points are denoted by red arrows.

Once skull reconstruction is complete, the user manually marks one unobstructed set of fiducial marks. Because the absolute spacing between these marks is known, the 3D skull mesh can then be rescaled to absolute dimensions.

The platform’s top surface is additionally equipped with a heating element to maintain the animal’s physiological temperature throughout surgery. Fixtures such as ear bars and bite bars (not shown) are integrated into the platform to secure the animal in place.

### 2.4 Automatic positioning of the animal’s skull to achieve the skull-flat position

Once the skull mesh has been reconstructed and scaled, the user manually marks the bregma and lambda points on the mesh within the 3D virtual space. The software calculates a virtual line connecting these two points, as well as a second line, perpendicular to the first, passing through its midpoint and running parallel to the two sides of the skull. Because the top surface of the Stewart platform is initially positioned parallel to the surgical table, the angle between these bregma-lambda-derived lines and the reference axes defined by the platform’s fiducial marks can be used to compute, via trigonometry, the angular offsets required to bring the skull into the skull-flat position. These positional adjustments are then displayed to the user for confirmation. Upon approval, the Stewart platform executes the adjustments, flattening the skull and preparing it for subsequent movement to the user-specified surgical coordinates.

### 2.5 Validation and quantification of error

To quantify the repeatability of the skull-flat position obtained by photogrammetric 3D reconstruction, a red laser pointer was affixed to a bare Mongolian gerbil (*Meriones unguiculatus*) skull, which was then mounted on the Stewart platform. The laser was aimed at the center of a sheet of graph paper positioned 265 mm from its tip, and the resulting laser-dot position was marked. A set of images was acquired of the skull using an iPhone 15 camera as described above, and a 3D mesh was generated using the algorithms detailed above. The platform was then moved to the skull-flat position calculated by the system, and the new laser-dot position was marked on the graph paper. The platform was then returned to its home position, and the entire cycle, image acquisition, mesh reconstruction, skull-flat positioning, and laser-dot marking, was repeated ten times. Centroid coordinates of the repeated laser-dot marks were extracted from scanned images of the graph paper using a custom Python-based image-analysis script. Positional deviations from the mean target location were converted into azimuth and elevation angular errors using the geometric relationship between lateral displacement and the arm length of the positioning system. Revisit accuracy was then summarized as the standard deviation and half-range of the angular errors across the ten repeated trials.

The system was further validated in vivo by targeting small, deep nuclei within the auditory brainstem of Mongolian gerbils. To this end, gerbils were anesthetized with isoflurane, and the scalp was reflected to expose the bregma and lambda landmarks. Skull-flat positioning was achieved using the photogrammetry-based method described above. Following skull-flat positioning, the medial nucleus of the trapezoid body (MNTB) and the lateral nucleus of the trapezoid body (LNTB) were targeted using coordinates obtained from a brain atlas (Radtke-Schuller et al. 2016). These nuclei are located very deep within the brainstem, bordering its ventral surface, making them challenging targets for traditional stereotaxic methods. In addition, these targets must be approached at an angle to avoid major blood vessels, since damaging them can cause excessive bleeding, further complicating surgery in this region. For MNTB, a craniotomy was drilled 4 mm caudal and 0.8 mm lateral to lambda. For LNTB, the craniotomy was drilled 4 mm caudal and 2.2 mm lateral to lambda. In both cases, the platform was tilted 20° forward from the skull-flat position. A glass pipette filled with the fluorescent tracer Dextran, Tetramethylrhodamine, and Biotin (3,000 MW; Thermo Fisher Scientific, Waltham, MA) was then advanced into the brain by 7500 µm (MNTB) or 7325 µm (LNTB). Gerbils were then euthanized and transcardially perfused with 4% paraformaldehyde (PFA). Brains were extracted, post-fixed in 4% PFA overnight, embedded in 4% agar, and sectioned on a vibratome (Leica VT 1000S, Wetzlar, Germany) at 100 to 200 µm. Sections were mounted in Fluoromount-G (Invitrogen, Carlsbad, CA) and imaged on an Olympus Fluoview FV1000 laser scanning confocal microscope (Tokyo, Japan).

## 3. Results

To evaluate the photogrammetry-based stereotaxic system, we first characterized the speed and quality of three-dimensional skull reconstruction, then quantified the repeatability of the resulting skull-flat positioning, and finally validated the system’s targeting accuracy in vivo by labeling small, deep nuclei in the auditory brainstem.

Figure 6(a) shows a representative set of images captured with an iPhone 15 camera, which were processed to generate the 3D mesh shown in figure 6(b). To evaluate the photogrammetry pipeline independently of biological variability, a 3D-printed rodent model with a Mongolian gerbil skull affixed to the head was used as the imaging target. Thirty images were captured from multiple angles around the model, following the acquisition protocol described above, and submitted to the photogrammetry reconstruction routine for mesh generation. Reconstruction was performed on a workstation equipped with an Intel Core i9-12900K processor, 48 GB of RAM, and an Nvidia GeForce RTX 4060 GPU, and required approximately 132 seconds from image upload to completed mesh. This processing time compares favorably with the time typically required to manually palpate and measure even a small number of skull landmarks, and it scales primarily with the number of input images rather than with the geometric complexity of the skull, making the approach practical for routine use within a surgical workflow.

The resulting mesh resolved the plastic head model, the affixed gerbil skull, the top plate of the Stewart platform, and the two ear bars used to secure the head, all within a single coordinate frame. Critically, the mesh was rendered in full color rather than as a uniform grayscale surface; across the exposed skull region in particular, natural variation in bone coloration and surface texture along the cranial sutures was clearly preserved. This stands in contrast to the monochromatic meshes produced by the structured-illumination method described above, in which the absence of color information made suture landmarks substantially harder to distinguish from the surrounding bone. The polychromatic rendering achieved by photogrammetry therefore directly addresses one of the principal limitations of the earlier system and was sufficient to permit clear, unambiguous identification of the bregma and lambda landmarks.

Once the photogrammetric reconstruction of the skull was complete, the bregma and lambda suture landmarks could be identified directly on the colored mesh. Figure 7(a) shows the manually marked three-dimensional coordinates of bregma and lambda on the reconstructed skull mesh. These two points were used to compute the anteroposterior tilt of the skull. In addition, the midpoint of the bregma-lambda line was calculated and used to define two corresponding lateral points on the skull, which together with bregma and lambda established a four-point reference plane from which both the lateral (roll) and yaw angles of the skull could be estimated (figure 7(b)).

Correcting yaw orientation as part of the automated skull-flattening procedure represents a meaningful methodological improvement over most existing stereotaxic systems, in which this adjustment, if performed at all, is typically done by visual estimation alone. An uncorrected yaw rotation leaves the skull’s mediolateral axis misaligned with the stereotaxic frame’s reference axes even when anteroposterior and lateral tilt have both been properly corrected. For targets located off the sagittal midline, particularly those approached at a non-zero insertion angle, this rotational misalignment introduces a systematic mediolateral targeting error that grows with both insertion depth and the lateral distance of the target from the rotation center. Because the present system derives yaw directly from the measured three-dimensional positions of the bregma and lambda landmarks and their associated lateral reference points, this correction is applied automatically and reproducibly, independent of operator judgment. The four-point reference plane defined by these landmarks was then compared mathematically to the reference plane defined by selected fiducial points on the Stewart platform surface (figure 7(a)), and the platform was moved accordingly until the skull-flat orientation was achieved.

To quantify the repeatability of the skull-flat position obtained using this method, a laser pointer was affixed to a gerbil skull mounted on the Stewart platform, and ten independent cycles of image acquisition, photogrammetric reconstruction, and skull-flattening were performed as described in the Methods. Across these ten trials, the azimuthal error was 0.00° ± 0.86° (S.D.; ±1.23° half-range), and the elevation error was −0.36° ± 0.72° (S.D.; ±1.27° half-range) (figure 8). The near-zero mean azimuthal error indicates no systematic directional bias in the lateral correction, while the small negative mean elevation error suggests a modest, consistent offset in the anteroposterior correction, distinct from random trial-to-trial variability, given that its standard deviation was comparable to that of the azimuthal measurement.

**Figure 8.**
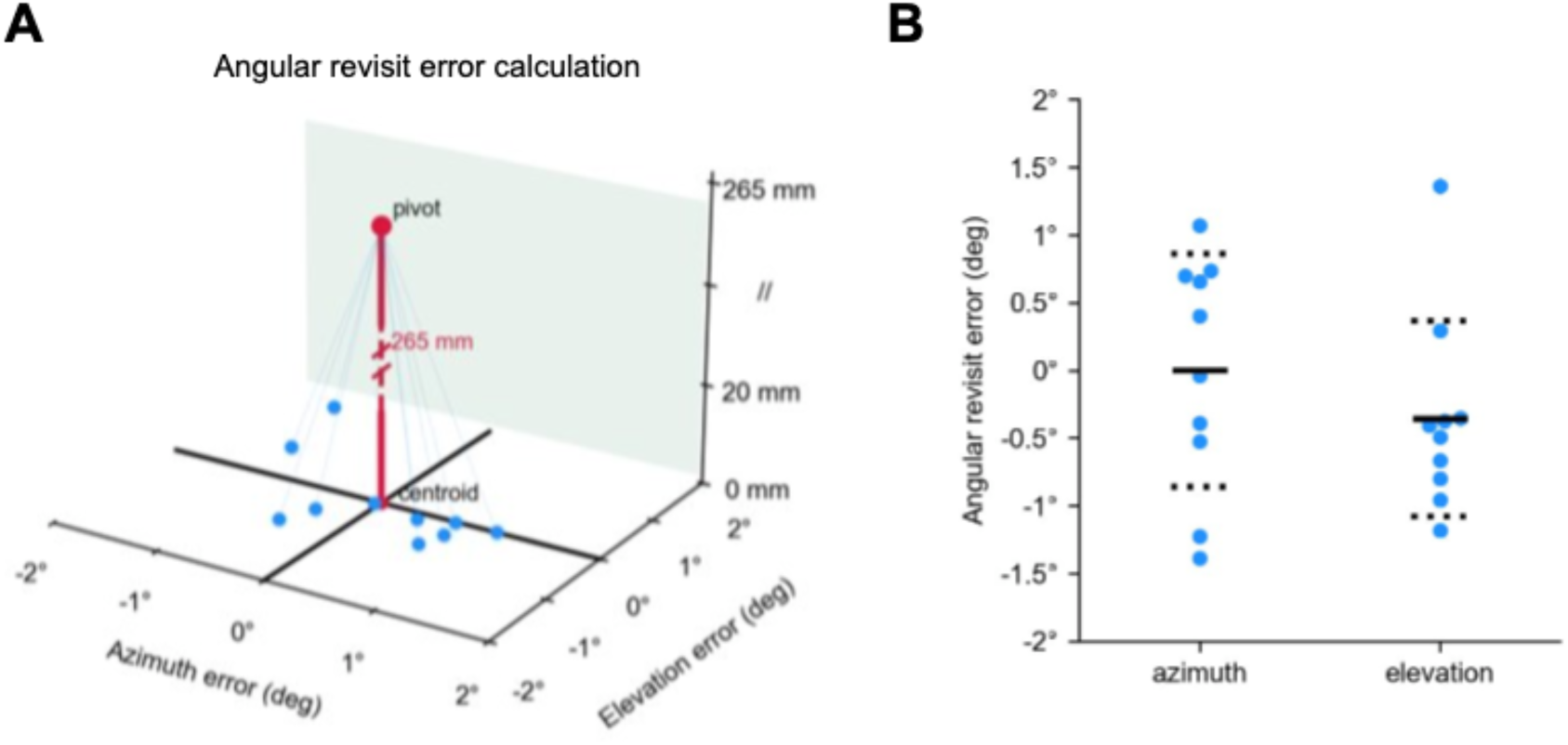
Angular repeatability error quantification. (A) Schematic of angular revisit error estimation based on repeated skull-flattening repositioning relative to a target centroid. Angular deviations were calculated from positional displacement relative to the pivot-to-target distance (265 mm). (B) Measured angular revisit errors across 10 repeated trials. Azimuth revisit error was 0.00° ± 0.86° S.D. (± 1.23 ° half-range), and elevation revisit error was −0.36° ± 0.72° S.D. (± 1.27 ° half-range). (blue dot: individual measurements; solid line: mean; dash line: ±1 S.D.)

Because angular positioning errors are amplified geometrically with insertion depth, even small angular deviations can translate into meaningful absolute targeting errors when reaching deep brain structures. As an approximate illustration, using the relationship that linear deviation is roughly equal to insertion depth multiplied by the tangent of the angular error, the measured half-range errors correspond to a worst-case combined positional deviation of approximately 230 micrometers at the insertion depths used to target the MNTB and LNTB in this study (7500 and 7325 micrometers, respectively). The true three-dimensional error for any given trajectory will depend on the specific combination of insertion angle and depth, but this estimate indicates that the angular accuracy achieved by the system is well suited to reliably reaching small, deeply situated nuclei, consistent with the in vivo results described below and the targeting outcomes shown in figure 9.

**Figure 9.**
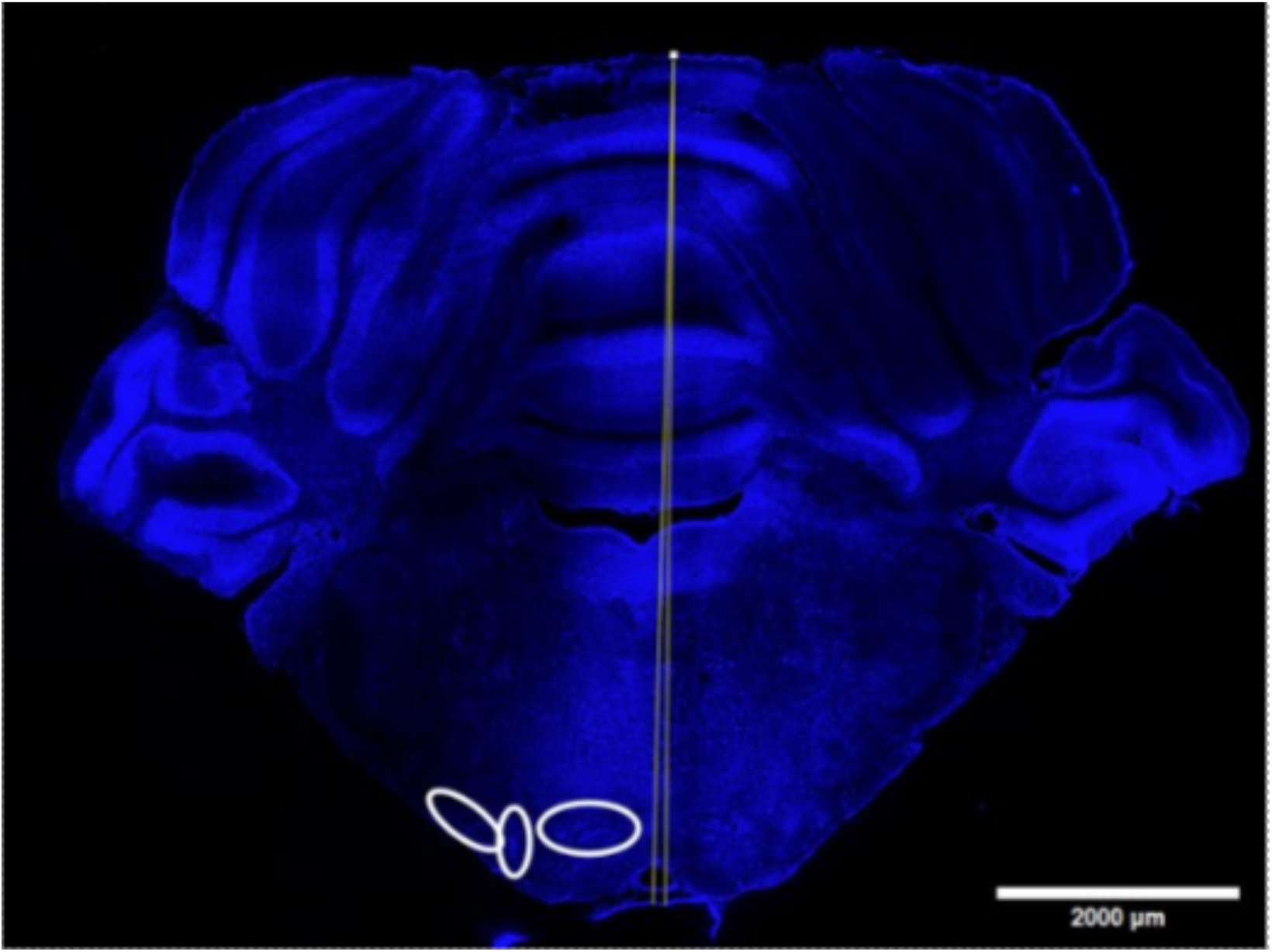
Inaccuracies caused by typical angular deviations. A coronal section of a Mongolian gerbil brain through the auditory brainstem. The yellow lines comprise a 0.85 degree angle, consistent with the standard deviation observed in repeated tests of the precision of the yaw measurement. White circles approximate the boundaries of nuclei in this region. Left to right: LNTB, MSO, and MNTB.

To assess the performance of the system under practical, in-vivo surgical conditions, fluorescent tracer was injected into the medial and lateral nuclei of the trapezoid body (MNTB and LNTB, respectively), two small nuclei situated deep within the auditory brainstem that are notoriously difficult to target using conventional stereotaxic methods, owing to their small size, their position near the ventral brainstem surface, and the need to approach them at an angle to avoid major blood vessels. Despite these challenges, both nuclei were successfully and consistently labeled following targeting based on the photogrammetry-derived skull-flat position and atlas coordinates (figure 10b and 10c). The reliability of tracer labeling across both nuclei indicates that the angular and positional accuracy quantified in the repeatability experiments above is sufficient to support precise targeting under real surgical conditions, including the angled trajectories required to avoid vasculature in this brainstem region, and not only under the idealized conditions of a static skull or phantom model.

**Figure 10.**
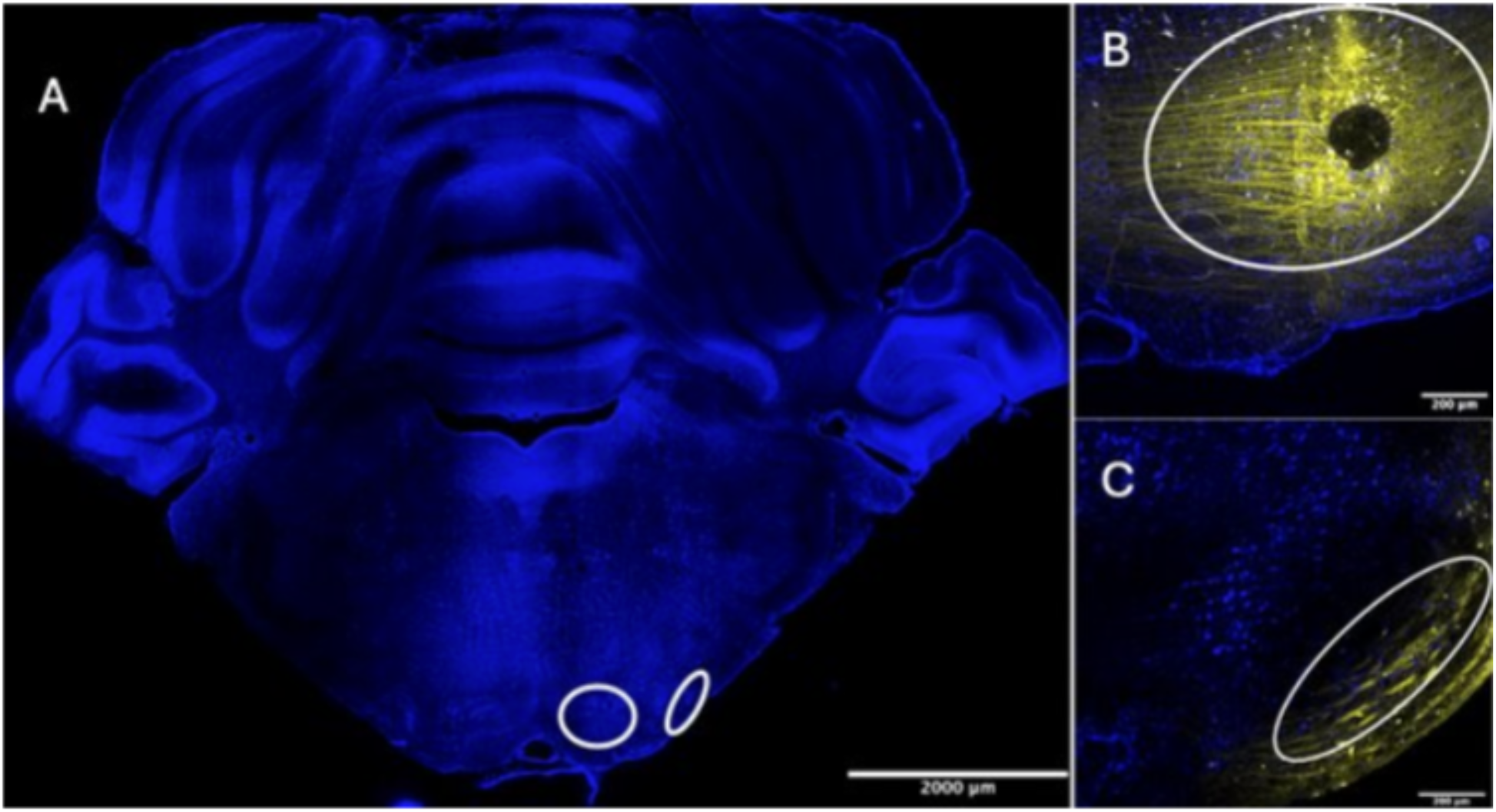
An example of a typical target accuracy accomplished with this technology. (A) A coronal slice through the superior olivary complex. White circles indicate the approximate boundary of MNTB (left) and LNTB (right). (B) A close-up image of MNTB with dextran tracer. The white circle indicates the approximate boundary of the nucleus. (C) A close-up image of LNTB with dextran tracer. The white circle indicates the approximate boundary of the nucleus.

## 4. Discussion

This study shows that replacing structured illumination with photogrammetry resolves the two principal limitations structured illumination based stereotaxic platforms, namely the bulky overhead projector and camera array and the difficulty of identifying suture landmarks on a monochromatic mesh. These improvements were accomplished without sacrificing targeting performance. The resulting system reconstructs a full color, micrometer scale skull surface from a brief series of handheld camera images, restores the visual cues a surgeon would normally use to find bregma and lambda, and still achieves sub-degree revisit precision. At the insertion depths used here, this corresponds to a worst-case positional deviation of roughly 230 micrometers, a margin narrow enough to reliably and repeatedly reach small and deep nuclei located ventrally in the brain stem, such as the nuclei MNTB and LNTB targeted here. Such brain areas are difficult to target because of their small size, their ventral location, and the angled trajectories needed to avoid major blood vessels. Taken together, these results suggest that color-resolved photogrammetry is not simply a convenience upgrade. It addresses a registration problem, accurate identification of cranial landmarks on an otherwise featureless surface, that had constrained the accuracy of the prior generation system.

The predecessor system (Ly et al., 2021) established that combining three-dimensional skull profiling with a 6DOF Stewart platform could improve stereotaxic accuracy and speed over manual methods, and the present work builds directly on that foundation. The advance here is not three-dimensional reconstruction itself, but the method used to obtain it. Structured illumination requires precise, fixed placement of a projector and a stereo camera pair, occupies space above the animal that may otherwise be needed for surgical tools. Structured illumination produces a mesh with no color information, so bregma and lambda must be inferred from subtle changes in surface geometry alone. Photogrammetry removes all three constraints at once. A single, freely handheld camera replaces the fixed optical hardware, the natural coloration of the skull is preserved in the reconstruction, and accuracy improved rather than degraded despite the simpler hardware. This indicates that, for this application, whatever spatial information is lost by abandoning a fixed, calibrated optical geometry is more than offset by the gain in landmark visibility and the larger number of viewing angles a freely moved camera can capture.

The system also compares informatively with current commercial stereotaxic platforms, although no head-to-head benchmarking against these instruments was performed in this study, and the comparisons below are drawn from published specifications rather than direct empirical testing. The Angle Two and Angle Three instruments (Leica Biosystems, Nussloch, Germany) use linear and rotary encoders to track manipulator position and apply mathematical corrections for skull tilt based on a small number of probed points; Angle Three extends this correction to yaw, estimated from any offset between bregma and lambda relative to the sagittal midline. Neurostar’s (Tübingen, Germany) Drill and Injection Robot applies a similar landmark based, atlas referenced correction for pitch, roll, and yaw, and additionally automates craniotomies within one workflow. The Model 71000 apparatus (RWD Life Science, Shenzhen, China) takes a different approach still, using atlas-based calibration referenced to bregma to position the manipulator without an explicit skull leveling step at all. Across all three platforms, skull orientation is inferred from a handful of discrete, manually probed points rather than from a continuous reconstruction of the skull surface. The present system instead derives pitch, roll, and yaw simultaneously from a four-point reference plane computed directly from a dense three-dimensional reconstruction of the entire exposed skull. This may offer a more spatially complete estimate of orientation, particularly for skulls with asymmetric or irregular suture geometry where a single bregma-lambda line could be less representative of the true mediolateral axis, though whether this yields a measurable accuracy advantage over these platforms remains to be tested directly.

Other academic groups have pursued automated, vision guided cranial surgery from different angles entirely. Craniobot (Ghanbari et al., 2019) couples a contact-based skull surface profiler with a computer numerically controlled milling system to automate the craniotomy itself, rather than skull flat positioning for deep injections or implants. The contrast between a contact-based profiler and the camera-based photogrammetry used here shows that there is more than one workable path toward removing manual measurement from cranial surgery, and that the better approach likely depends on the procedural goal: automating the craniotomy versus automating full six axis skull alignment for deep targeting.

Several limitations should be considered when interpreting these results. Bregma and lambda are still marked manually on the reconstructed mesh, so operator judgment, now aided substantially by color information, remains part of the workflow. Fully automating landmark detection would remove this remaining source of variability. Mesh reconstruction required approximately 132 seconds in this study, fast relative to manual measurement but not instantaneous, and image quality depends on adherence to the acquisition protocol described in the Methods, including consistent lighting and zoom across the image set. The small but consistent negative elevation error (−0.36°), distinct from the near zero azimuthal error, suggests a residual systematic offset in the anteroposterior calibration that merits further investigation rather than dismissal as trial-to-trial noise. As noted above, the comparisons drawn with commercial platforms in this discussion rest on published specifications rather than controlled, side by side testing, and such a comparison would be required before any quantitative claim of relative advantage could be substantiated.

These limitations point toward several directions for future work. Automated, learning based detection of bregma and lambda directly from the colored mesh could remove the one manually measured step remaining in the workflow. Reducing reconstruction time, whether through algorithmic optimization, dedicated hardware, or a shift toward continuous video-based capture, would further streamline integration into routine surgical use. Extending validation to additional species, ages, and brain targets, and conducting direct comparative studies against existing commercial platforms, would help establish the generality of the approach and clarify where it offers the greatest practical benefit.

Overall, these findings indicate that color resolved photogrammetry, combined with a 6DOF platform capable of automated pitch, roll, and yaw correction, offers a practical solution to a registration limitation that persists across current stereotaxic platforms, both commercial and academic: achieving accurate, repeatable, full skull alignment without dedicated overhead optical hardware or reliance on a small number of manually probed points. This is likely to matter most for small, deep, and laterally displaced targets such as MNTB and LNTB, where even modest angular error is geometrically amplified at depth, but the underlying approach is general and should extend to a broad range of stereotaxic targets and species.

## Acknowledgement & Funding

Supported by NIH R41 NS 119079 to PopNeuron LLC

